# Domain-relevant auditory expertise modulates the additivity of neural mismatch responses in humans

**DOI:** 10.1101/541037

**Authors:** Niels Chr. Hansen, Andreas Højlund, Cecilie Møller, Marcus Pearce, Peter Vuust

## Abstract

It is unknown whether domain-relevant expertise is associated with more independent or more dependent predictive processing of acoustic features. Here, mismatch negativity (MMNm) was recorded with magnetoencephalography (MEG) from 25 musicians and 25 non-musicians, exposed to complex musical multi-feature and simple oddball control paradigms. Deviants differed in frequency (F), intensity (I), perceived location (L), or any combination of these (FI, IL, LF, FIL). Neural processing overlap was assessed through MMNm additivity by comparing double- and triple-deviant MMNms (“empirical”) to summed constituent single-deviant MMNms (“modelled”). Significantly greater subadditivity was present in musicians compared to non-musicians, specifically for frequency-related deviants in complex contexts. Despite using identical sounds, expertise effects were absent from the simple paradigm. This novel finding supports the *dependent processing hypothesis* whereby experts recruit overlapping neural resources facilitating more integrative representations of domain-relevant stimuli. Such specialized predictive processing may enable experts such as musicians to capitalise on complex acoustic cues.

## Introduction

The ability to distinguish and combine features of sensory input guides behaviour by enabling humans to engage successfully with perceptual stimuli in their environment (Crosse, Butler, and Lalor 2015; Hommel 2004). While sophisticated models exist for visual feature processing (Treisman and Gelade 1980; Nassi and Callaway 2009; Di Lollo 2012; Grill-Spector and Weiner 2014), auditory objects transform over time and remain more elusive (Griffiths and Warren 2004; Shamma 2008). Modality-specific divergences may therefore be expected. Indeed, findings that auditory feature conjunctions are processed pre-attentively (Winkler, Czigler, Sussman, Horváth and Balázs 2005) and faster than single features (Woods, Alain and Ogawa 1998) and that the identity features pitch and timbre sometimes take precedence over location (Maybery et al. 2009; Delogu, Gravina, Nijboer and Postma 2014) deviate from findings in visual perception (Campo et al. 2010; but see Guérard, Morey, Lagacé and Tremblay 2013). Although auditory feature integration mechanisms are partly congenital (Ruusuvirta 2001; Ruusuvirta, Huotilainen, Fellman and Näätänen 2003) and subject to evolutionary adaptation (Fay and Popper 2000), it remains unknown whether they vary with auditory expertise levels. Given its early onset and persistence throughout life (Ericsson 2006), musicianship offers an informative model of auditory expertise (Vuust et al. 2005; Stewart 2008; Herdener et al. 2010; Herholz and Zatorre 2012; Schlaug 2015).

Electroencephalography (EEG) and magnetoencephalography (MEG) enable quantification of feature integration using the additivity of the mismatch negativity response (MMN) and its magnetic counterpart (MMNm), respectively. The MMN(m) itself represents a deflection in the event-related potential or field (ERP/ERF) peaking around 150-250 ms after presentation of an unexpected stimulus (Näätänen, Gaillard, & Mäntysalo 1978). It results from active cortical prediction rather than from passive synaptic habituation (Wacongne, Changeux and Dehaene 2012). By comparing *empirical* MMN(m)s to double or triple deviants (differing from standards on two or three features) to *modelled* MMN(m)s obtained by summing the MMN(m) responses for the constituent single deviants, inferences have been made about the potential overlap in neural processing (e.g., Levänen, Hari, McEvoy and Sams 1993). Correspondence between empirical and modelled MMN(m)s is interpreted to indicate independent processing whereas subadditivity—where modelled MMN(m)s exceed empirical MMN(m)s—suggests overlapping, dependent processing.

Using MMN(m) additivity, segregated feature processing has been estabnlished for frequency, intensity, onset asynchrony, and duration (Levänen, Hari, McEvoy and Sams 1993; Schröger 1995; Paavilainen, Valppu and Näätänen 2001; Wolff and Schröger 2001; Paavilainen, Mikkonen, et al. 2003) with spatially separate neural sources (Giard et al. 1995; Levänen, Ahonen, Hari, McEvoy and Sams 1996; Rosburg 2003; Molholm, Martinez, Ritter, Javitt and Foxe 2005). MMN(m) is also additive for inter-aural time and intensity differences (Schröger 1996), phoneme quality and quantity (Ylinen, Huotilainen and Näätänen 2005), and attack time and even-harmonic timbral attenuation (Caclin et al. 2006). Feature conjunctions occurring infrequently in the local context, moreover, result in distinct MMN(m) responses (Gomes, Bernstein, Ritter, Vaughan and Miller 1997; Sussman, Gomes, Nousak, Ritter and Vaughan 1998) that are separable in terms of neural sources (Takegata, Huotilainen, Rinne, Näätänen and Winkler 2001) and extent of additivity from those elicited by constituent (Takegata, Paavilainen, Näätänen and Winkler 1999) or more abstract pattern deviants (Takegata, Paavilainen, et al. 2001).

Conversely, subadditive MMN(m)s occur for direction of frequency and intensity changes (Paavilainen, Degerman, Takegata and Winkler 2003), for aspects of timbre (Caclin et al. 2006), and between frequency deviants in sung stimuli and vowels (Lidji, Jolicœur, Moreau, Kolinsky and Peretz 2009) and consonants (Gao et al. 2012). Generally, subadditivity is greater for triple than double deviants (Paavilainen et al. 2001; Caclin et al. 2006). While it is known that expertise modulates MMN(m) responses (Koelsch, Schröger and Tervaniemi 1999; Vuust, Ostergaard, Pallesen, Bailey and Roepstorff 2009), it remains unknown whether it modulates MMN(m) additivity.

The current MEG study aims to investigate whether MMNm additivity varies as a function of musical expertise. Specifically, two hypotheses are contrasted. First, the *independent processing hypothesis* posits that expertise is associated with specialised feature processing by separate neural populations, increasing access to lower-level representations that have higher context-specific relevance (Ahissar and Hochstein 2004; Ahissar, Nahum, Nelken and Hochstein 2009). Second, the *dependent processing hypothesis* posits that expertise is associated with processing of multiple features by shared neural resources, manifesting as decreased neural activity (Jäncke, Gaab, Wüstenberg, Scheich and Heinze 2001; Zatorre, Delhommeau and Zarate 2012). Since expertise produces more accurate expectations (Hansen and Pearce 2014; Hansen, Vuust and Pearce 2016) and shorter MMN(m) latencies (Lappe, Lappe and Pantev 2016) in musical contexts specifically, these hypotheses will be tested using contrasting paradigms with higher and lower levels of complexity and corresponding musical relevance.

## Results

To anticipate our results, significantly greater MMNm subadditivity was found in musicians compared to non-musicians for frequency-related features in the complex musical multi-feature paradigm. These expertise effects were absent from the simple control paradigm. Further details are reported separately for the two paradigms below.

### Complex musical multi-feature paradigm

In the main experimental paradigm—the complex musical multi-feature paradigm—all seven single, double, and triple deviants elicited significantly larger responses than the standards for musicians as well as for non-musicians (Table 1), indicative of significant MMNm responses in all conditions of this paradigm. Moreover, the triple deviant resulted in a significantly larger MMNm than each of the double deviants (Table 2). Except for a single comparison between the location deviant and the double deviant combining location and frequency, MMNms to all double deviants were significantly larger than MMNms to single deviants, indicating that the addition of an extra feature significantly increased the MMNm amplitude.

**Table 1.**
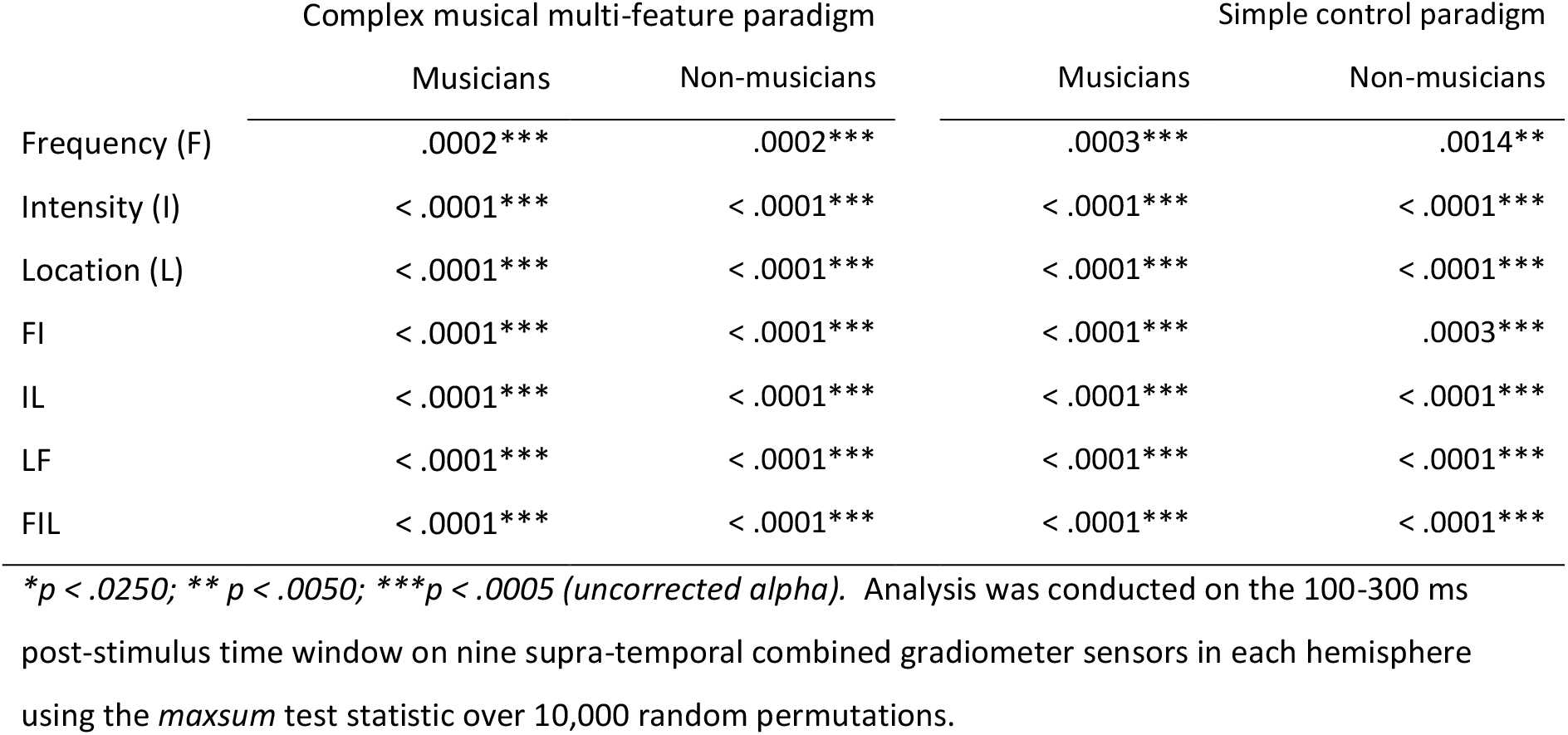
Magnetic mismatch negativity response (MMNm) for single deviants in Frequency (F), Intensity (I), Location (L), as well as double and triple deviants combining deviants on these features (FI, IL, LF, FIL). Monte Carlo *p* values from non-parametric, cluster-based permutation tests of deviant > standard.

|  | Complex musical multi-feature paradigm |  | Simple control paradigm |  |
| --- | --- | --- | --- | --- |
|  | Musicians | Non-musicians | Musicians | Non-musicians |
| Frequency (F) | .0002*** | .0002*** | .0003*** | .0014** |
| Intensity (I) | < .0001*** | < .0001*** | < .0001*** | < .0001*** |
| Location (L) | < .0001*** | < .0001*** | < .0001*** | < .0001*** |
| FI | < .0001*** | < .0001*** | < .0001*** | .0003*** |
| IL | < .0001*** | < .0001*** | < .0001*** | < .0001*** |
| LF | < .0001*** | < .0001*** | < .0001*** | < .0001*** |
| FIL | < .0001*** | < .0001*** | < .0001*** | < .0001*** |

\* $p < .0250$ ; \*\* $p < .0050$ ; \*\*\* $p < .0005$ (uncorrected alpha). Analysis was conducted on the 100-300 ms post-stimulus time window on nine supra-temporal combined gradiometer sensors in each hemisphere using the *maxsum* test statistic over 10,000 random permutations.

**Table 2.** Potential additivity of the MMNm response, as evident from comparisons of triple (FIL) with double deviants (FI, IL, LF) and double with single deviants (F, I, L). Monte Carlo *p* values from non-parametric, cluster-based permutation tests of triple > double > single deviants.

|  | Complex musical multi-feature paradigm | Simple control paradigm |
| --- | --- | --- |
| FIL > FI | < .0001*** | < .0001*** |
| FIL > IL | .0033** | .0003*** |
| FIL > LF | < .0001*** | < .0001*** |
| FI > F | < .0001*** | .0003*** |
| FI > I | .0203* | .0005*** |
| IL > I | < .0001*** | < .0001*** |
| IL > L | < .0001*** | < .0001*** |
| LF > L | n.s. | .0002*** |
| LF > F | < .0001*** | .0093* |

F: Frequency; I: Intensity; L: Location; \* $p < .0250$ ; \*\* $p < .0050$ ; \*\*\* $p < .0005$ (*uncorrected alpha*). Analysis was conducted on the 100-300 ms post-stimulus time window on nine supratemporal combined gradiometer sensors in each hemisphere using the *maxsum* test statistic over 10,000 random permutations.

The non-significant LF vs. L comparison as well as the somewhat larger *p* values for the comparisons between responses elicited with and without the frequency component (i.e. FIL vs. IL, FI vs. I) already indicated that a certain extent of subadditivity was present specifically for the frequency component. This is also suggested by the dissimilarity between modelled and empirical responses in the first, third, and fourth rows of Figure 1. The subsequent main analysis addressed how this possible effect interacted with musical expertise.

**Fig. 1.**
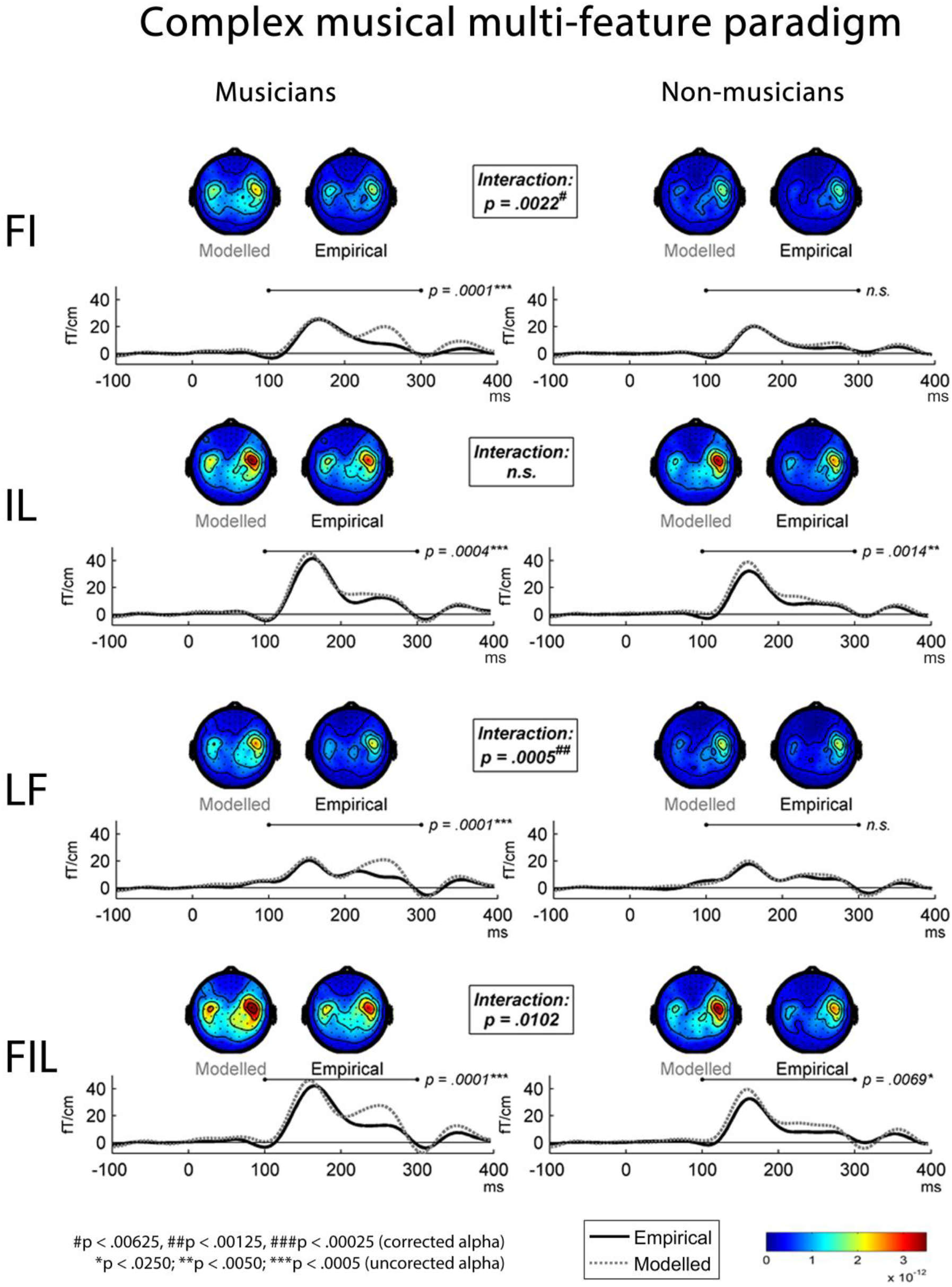
Modelled and empirical mismatch negativity (MMNm) in musicians (*n* = 25) and non-musicians (*n* = 25) in the complex musical multi-feature paradigm in response to deviants differing in either frequency & intensity (FI), intensity & location (IL), location & frequency (LF), or frequency & intensity & location (FIL). Modelled MMNms (dashed grey lines) correspond to the sum of the MMNms to two or three single deviants whereas empirical MMNms (solid black lines) correspond to MMNms obtained for double and triple deviants. Comparing the plots for modelled and empirical MMNms, cluster-based permutation tests showed significantly greater subadditivity in musicians compared to non-musicians specifically in the double deviants involving frequency (i.e. FI and LF) (Table 3). This expertise-by-additivity effect was marginally non-significant for the triple deviant (i.e. FIL). The event-related field (ERF) plots depict the mean of the data from the peak sensor and eight surrounding sensors in each hemisphere (the peak gradiometer pairs were MEG1342+1343 (right) and MEG0242+0243 (left)). Low-pass filtering at 20 Hz was applied for visualisation purposes only. The topographical distributions depict the 100-300 ms post-stimulus time range for the difference waves between standard and deviant responses, and *p* values outside the boxes reflect simple effects of additivity separately for musicians and non-musicians.

Indeed, the differences between modelled and empirical responses were significantly greater in musicians than in non-musicians for the FI and LF deviants (Table 3). For the triple deviant (FIL), this interaction effect approached significance whereas it was absent for the IL deviant, which did not include the frequency component. Consistent with this picture, follow-up analyses of the significant interactions found simple additivity effects only for musicians (Table 3, Figure 1).

**Table 3.**
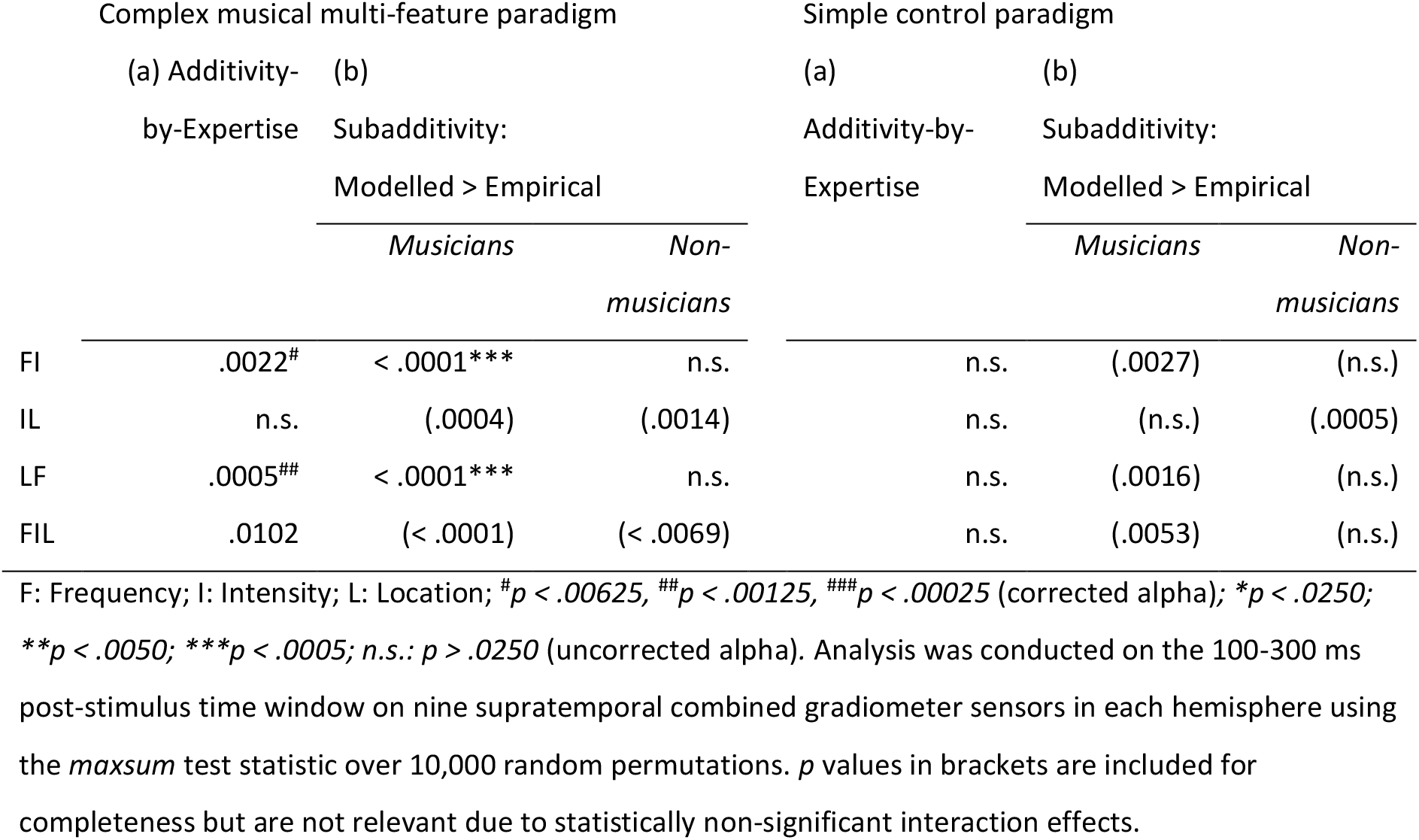
Subadditivity of the MMNm response. Monte Carlo *p* values from non-parametric, cluster-based permutation tests of (a) the interaction of additivity-by-expertise, and (b) simple effects of additivity for musicians and non-musicians.

### Simple control paradigm

In the simple control paradigm, significant MMNm effects were also present for all deviant types (Table 1). Here, additivity was more uniformly present in terms of significant differences between all single and double deviants as well as between the double deviants and the triple deviant (Table 2). The main analysis showed no differences in additivity between the two groups and across the various deviant types (Table 3). This pattern also emerges from the plots of event-related fields and scalp topographies in Figure 2.

**Fig. 2.**
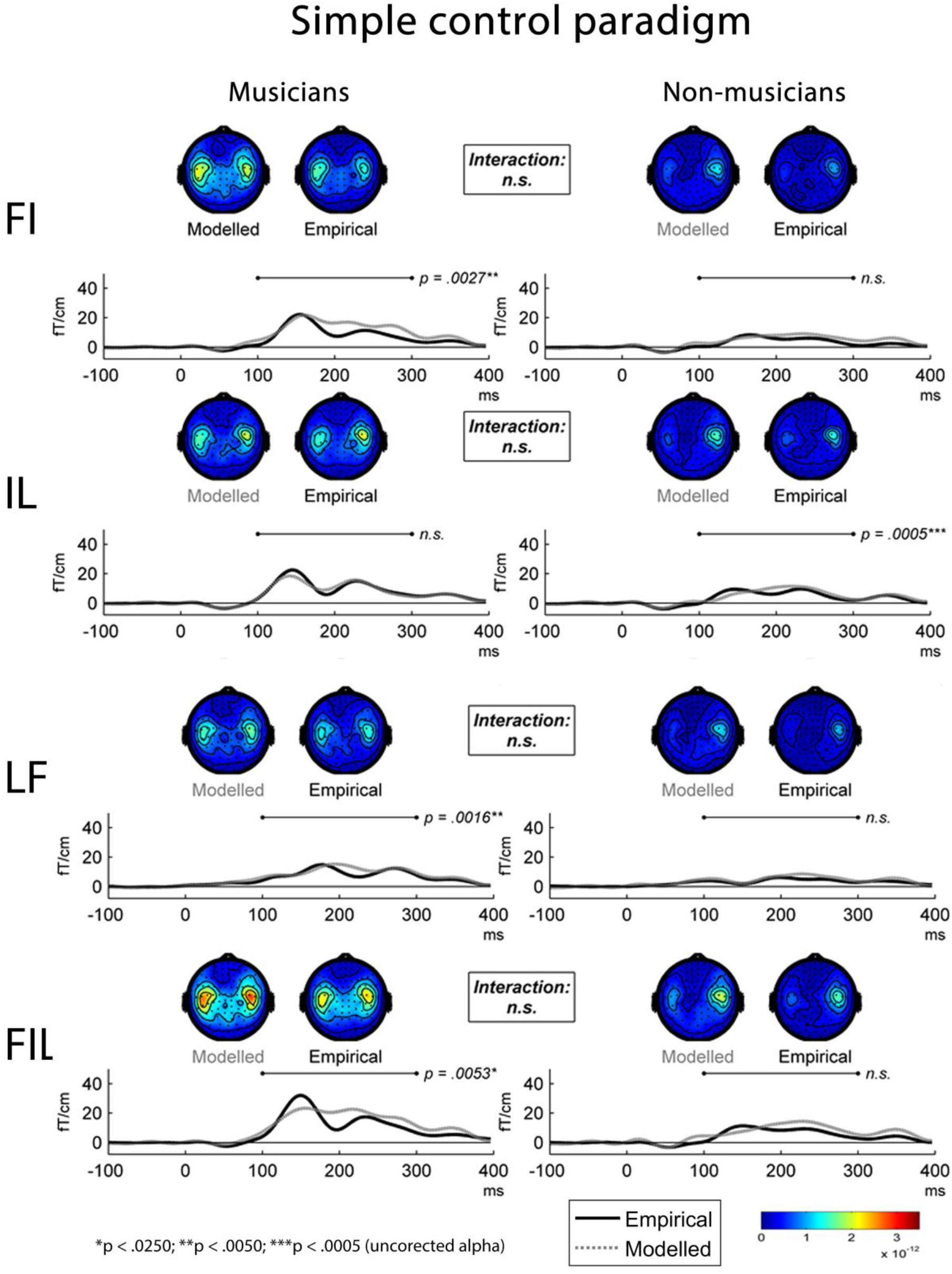
Modelled and empirical mismatch negativity (MMNm) in in musicians (*n* = 25) and non-musicians (*n* = 25) in the simple control paradigm in response to double deviants differing in either frequency & intensity (FI), intensity & location (IL), or location & frequency (LF), as well as to triple deviants differing in frequency & intensity & location (FIL). Modelled MMNms (dashed grey lines) correspond to the sum of MMNms to two or three single deviants whereas empirical MMNms (solid black lines) correspond to the actual MMNms obtained with double and triple deviants. In contrast to the musical multi-feature paradigm, cluster-based permutation tests showed no significant differences in additivity between musicians and non-musicians when comparing the modelled and empirical MMNms (Table 3). The event-related field (ERF) plots depict the mean of the data from the peak sensor and eight surrounding sensors in each hemisphere (the peak gradiometer pairs were MEG1342+1343 (right) and MEG0242+0243 (left)). Low-pass filtering at 20 Hz was applied for visualisation purposes only. The topographical distributions depict the 100-300 ms post-stimulus time range for the difference waves between standard and deviant responses, and *p* values reflect simple effects of additivity separately for musicians and non-musicians.

## Discussion

### Expertise as dependent feature processing

The current results provide the first neurophysiological evidence of more dependent processing of multiple auditory feature deviants in musical experts compared to non-experts. In a complex musical paradigm, magnetic mismatch negativity responses (MMNm) elicited by double deviants differing in multiple features (including frequency) were lower than the sum of MMNm responses to the constituent single-feature deviants. This selective subadditivity for the musically relevant frequency feature was either absent or present to a smaller extent in non-musicians. While relatively small MMNm amplitudes in non-musicians may be suggestive of floor effects, a significant MMNm was found in all conditions. Therefore, statistical analysis should have revealed any true subadditivity in non-experts. In addition to the main finding of expertise-related differences in frequency-related subadditivity, the pattern of results between the two paradigms suggests that these differences were context-specific. Specifically, the expertise differences were absent when using identical sounds in a simpler and less musically-relevant configuration as a control paradigm.

We interpret these findings as support for the *dependent processing hypothesis* by which musical expertise is associated with enhanced processing by recruiting shared neural resources for more complex representations of domain-relevant stimuli. Importantly, our cross-sectional study design does not allow us to determine whether musical training causally induces dependent processing or whether pre-existing dependent processing benefits musicianship. A causal interpretation would contrast with formulations of feature integration theory regarding feature processing as innate or acquired through normal neurodevelopment and therefore largely immutable (Treisman and Gelade 1980; Quinlan 2003). Findings that perceptual learning modulates attention, thus determining whether specific features are considered for binding (Colzato, Raffone and Hommel 2006), have already questioned this view. Indeed, Neuhaus and Knösche (2008) demonstrated more integrative pitch and rhythm processing in musicians than non-musicians manifested in expertise-related differences in the P1 and P2 components. By extending these findings to intensity and location and relating them to MMNm responses, we consolidate the association of musical training with changes in predictive neural processing (Skoe and Kraus 2012).

Evidence for expertise-related effects on multimodal integration of audiovisual and audiomotor stimuli has steadily accumulated (Paraskevopoulos, Kuchenbuch, Herholz and Pantev 2012, 2014; Paraskevopoulos and Herholz 2013; Bishop and Goebl 2014; Proverbio, Calbi, Manfredi and Zani 2014; Pantev, Paraskevopoulos, Kuchenbuch, Lu and Herholz 2015; Proverbio, Attardo, Cozzi and Zani 2015; Møller et al. 2018). While integration *across* modalities appears less prominent in experts who may be better at segregating auditory features from audiovisual compounds (Møller et al. 2018), our study suggests that feature integration *within* the relevant modality may simultaneously increase with expertise.

Consistent with a causal interpretation of our data in the context of the *dependent processing hypothesis*, decreased neural activity is observed for perceptual learning in audition (Jäncke et al. 2001; Berkowitz and Ansari 2010; Zatorre et al. 2012) and vision (Schiltz et al. 1999; van Turennout, Ellmore and Martin 2000; Kourtzi, Betts, Sarkheil and Welchman 2005; Yotsumoto, Watanabe and Sasaki 2008). These changes are sometimes associated with enhanced effective connectivity (Büchel, Coull and Friston 1999), which may enable musicians to rely more heavily on auditory than visual information in audiovisual tasks (Møller et al. 2018; Paraskevopoulos, Kraneburg, Herholz, Bamidis and Pantev 2015). This seems adaptive assuming that auditory information is better integrated and thus potentially more informative and relevant to musicians.

More integrative auditory processing may indeed benefit musicians behaviourally. For instance, this may manifest in terms of enhanced verbal and visual memory (Chan, Ho and Cheung 1998; Jakobson, Lewycky and Kilgour 2008). This would be consistent with feature-based theories of visual short-term memory positing that integrative processing allows experts to incorporate multiple features into object representations, thus improving discrimination of highly similar exemplars (Curby, Glazek and Gauthier 2009).

### Frequency-related feature selectivity

Whereas MMNm subadditivity was present for double and triple deviants comprising frequency, intensity, and location, expertise only interacted with additivity for frequency-related deviants. This aligns well with the different levels of musical relevance embodied by these features, with frequency, intensity, and location representing decreasingly common parameters of syntactic organisation in music. More specifically, frequency is usually predetermined by composers in terms of unambiguous notation of intended pitch categories, the spontaneous changing of which would produce distinctly different melodies. Intensity is subject to more flexible expressive performance decisions (Palmer 1996; Widmer and Goebl 2004). Changing it would usually not compromise syntax or melodic identity. Sound source localization is equally crucial in the everyday lives of musicians and non-musicians, with the possible exception orchestral and choral conductors whose more advanced sound localization skills are evident from specialised neuroplasticity (Münte, Kohlmetz, Nager and Altenmüller 2001; Nager, Kohlmetz, Altenmuller, Rodriguez-Fornells and Münte 2003). Given that our musician group, however, only comprised a single individual with conducting experience, we can assume that all members of this group had intensive pitch-related training, that most members regularly used intensity as an expressive means, but that sound source localization was merely of secondary importance.

The prominence of pitch over intensity and location may relate to the diverging potential for learning effects to emerge within these features. For example, while learning of frequency discrimination is ubiquitous (Kishon-Rabin, Amir, Vexler and Zaltz 2001), adaptation to altered sound-localization cues is only partial in the horizontal plane (Hofman, Vlaming, Termeer and Van Opstal 2002; Wright and Zhang 2006). Indeed, recognising and producing pitch accurately is rehearsed intensively in musical practice (Besson, Schön, Moreno, Santos and Magne 2007). Remarkably, voice pedagogues regard pitch intonation as *the* most important factor in determining singing talent (Watts, Barnes-Burroughs, Andrianopoulos and Carr 2002).

One possible interpretation following from the present study is that musical learning may involve changes in feature integration which, accordingly, would manifest most clearly in the musically relevant and perceptually plastic pitch-related domain. Indeed, musical pitch information has been shown to be integrated with intensity (Grau and Kemler-Nelson 1988; McBeath and Neuhoff 2002) and timbre (Allen and Oxenham 2014; Hall, Pastore, Acker and Huang 2000; Krumhansl and Iverson 1992; Lidji et al. 2009), leading to nearly complete spatial overlap in neural processing of frequency and timbre (Allen, Burton, Olman and Oxenham 2017). More broadly, the three features included here exemplify both the “what” (frequency, intensity) and “where” (location, intensity) streams of auditory processing, which have been found to follow different patterns of feature integration (Delogu et al. 2014). By mostly recruiting participants with uniform expertise levels, many previous studies have avoided systematic group comparisons capable of demonstrating expertise-related differences in feature integration.

In sum, although frequency, intensity, and location all have relevance in music, frequency represents the most salient cue for identifying and distinguishing musical pieces and their component themes and motifs. Therefore, when musicians show selectively affected frequency processing, we infer this to be more likely due to their intensive training than to innate predispositions (Herdener et al. 2010; Hyde et al. 2009). Even if aspects of innately enhanced frequency-specific processing are present, they are likely to require reinforcement through intensive training, possibly motivated by greater success experiences as a young musician. While these learning effects may transfer to prosodic production and decoding (Besson et al. 2007; Pastuszek-Lipińska 2008; Lima and Castro 2011), they are of limited use outside musical contexts where sudden changes in intensity or location are usually more likely than frequency to constitute environmentally relevant cues.

### Context-dependency of expert feature processing

It is plausible that because superior frequency processing is merely adaptive in relevant contexts, musicians showed greater frequency-related feature integration for more complex musically relevant stimuli. This corresponds with stronger MMNm amplitudes, shorter MMNm latencies, and more widespread cortical activation observed in musically complex compared to simple oddball paradigms (Lappe, Lappe and Pantev 2016). Greater MMNm amplitudes and shorter latencies similarly emerge for spectrally rich tones (e.g., piano) compared to pure tones that only rarely occur naturally (Tervaniemi et al. 2000). Musical expertise, moreover, produces greater MMN amplitudes (Fujioka, Trainor, Ross, Kakigi and Pantev 2004) and more widespread activation (Pantev et al. 1998) for complex than for pure tones. Enhanced processing of audiovisual asynchronies in musicians is also restricted to music-related tasks, remaining absent for language (Lee and Noppeney 2014). Behaviourally, musical stimuli activate more specific predictions in musicians and thus evoke stronger reactions when expectations are violated (Hansen and Pearce 2014; Hansen, Vuust and Pearce 2016). Research in other expertise domains find comparable levels of domain-specificity (de Groot 1978; Ericsson and Towne 2010). Observations that pitch and duration in melodies are integrated when these dimensions covary— but not when they contrast—and that such integration increases with exposure (Boltz 1999) suggest an interpretation of our results where musicians are predisposed to capitalise more optimally on contextual cues. Note, though, that the two present paradigms did not only differ on complexity and musical relevance, but also on stimulus onset asynchrony, predictability of deviant location, and general textural complexity. Although these differences between the two paradigms do not compromise our main finding of expertise-related subadditivity for frequency-related features, future research should try to disentangle the individual contribution of these factors.

### Future research and unresolved issues

It is worth noting that increased frequency-related feature integration in musicians observed here seems inconsistent with other behavioural and neuroimaging results. For example, a behavioural study found that interference of pitch and timbre was unaffected by expertise (Allen and Oxenham 2014). Conversely, findings that pitch and consonants in speech produce subadditive MMN responses (Gao et al. 2012), but do not interact behaviourally (Kolinsky, Lidji, Peretz, Besson and Morais 2009), suggest that measures of behavioural and neural integration do not always correspond (Musacchia, Sams, Skoe and Kraus 2007). Additionally, cross-modal integration often results in increased rather than decreased neural activity (Stein 2012; Crosse, Butler and Lalor 2015). For instance, trumpeters’ responses to sounds and tactile lip stimulation exceed the sum of constituent unimodal responses (Schulz, Ross and Pantev 2003). When reading musical notation, musicians dissociate processing of spatial pitch in the dorsal visual stream from temporal movement preparation in the ventral visual stream (Bengtsson and Ullén 2006) and show no neurophysiological or behavioural interference of frequency and duration (Schön and Besson 2002). This suggests greater independence in experts.

The Reverse Hierarchy Theory of perceptual learning supports this view by asserting that expertise entails more segregated representations of ecologically relevant stimuli, thus providing access to more detailed lower-level representations (Ahissar and Hochstein 2004; Ahissar et al. 2009). This has some bearing on musical intuitions. Specifically, segregated processing would presumably better enable musicians to capitalise on the covariance of acoustic features, as demonstrated for tonal and metrical hierarchies (Prince and Schmuckler 2014). Phenomenal accents in music are indeed intensified by coinciding cues (Lerdahl and Jackendoff 1983; in review by Hansen 2011), which may underlie musicians’ superiority in decoding emotion from speech (Lima and Castro 2011) and segmenting speech (François, Chobert, Besson and Schön 2013; François, Jaillet, Takerkart and Schön 2014) and music (Deliège 1987; Peebles 2011, 2012). Given that individuals with better frequency discrimination thresholds more capably segregate auditory features from audiovisual stimuli (Møller et al. 2018), it remains unknown whether musicians suppress frequency-related feature integration whenever constituent features are irrelevant to the task at hand. The interaction of expertise with frequency-related feature-selectivity and context-specificity observed here contributes crucial empirical data to ongoing discussions of seemingly diverging findings of dependent and independent musical feature processing (Boltz 1999; Palmer and Krumhansl 1987; Tillmann and Lebrun-Guillaud 2006; Waters and Underwood 1999).

## Materials and Methods

### Participants

Twenty-five non-musicians (11 females; mean age: 24.7 years) and 25 musicians (10 females; mean age: 25.0 years) were recruited through the local study participation system and posters at Aarhus University and The Royal Academy of Music Aarhus. Members of the musician group were full-time conservatory students or professional musicians receiving their main income from performing and/or teaching music. The non-musician group had no regular experience playing a musical instrument and had received less than one year of musical training beyond mandatory music lessons in school. As shown in Table 4, the two groups were matched on age and sex, and musicians scored significantly higher on all subscales of *Goldsmiths Musical Sophistication Index*, v. 1.0 (Müllensiefen, Gingras, Musil and Stewart 2014).

**Table 4.** Demographics and musical sophistication of research participants.

|  | Musicians<br>(*n* = 25) | Non-musicians<br>(*n* = 25) | Chi-square test |  |
| --- | --- | --- | --- | --- |
| | | | $\chi^2$ | *p* |
| Gender | 10F, 15M | 11F, 14M | .082 | .774 |
|  | mean (SD) | mean (SD) | Mann-Whitney test |  |
|  |  |  | *U* | *p* |
| Age | 25.0 (3.9) | 24.7 (2.9) | 302.5 | .845 |
| GMSI1: Active engagement | 49.7 (5.3) | 28.3 (10.6) | 26.5 | < .001 |
| GMSI2: Perceptual abilities | 55.3 (4.5) | 37.7 (6.7) | 12.5 | < .001 |
| GMSI3: Musical training | 42.4 (3.1) | 10.9 (3.2) | 0.0 | < .001 |
| GMSI4: Emotions | 36.2 (4.0) | 27.7 (5.0) | 61.0 | < .001 |
| GMSI5: Singing abilities | 38.9 (5.1) | 21.2 (5.7) | 4.0 | < .001 |
| GMSI6: General sophistication | 104.9 (7.3) | 49.4 (11.1) | 0.0 | < .001 |

GMSI1-6 designate the six subscales of *Goldsmiths Musical Sophistication Index* v1.0 (Müllensiefen et al. 2014).

All participants were right-handed with no history of hearing difficulties. Informed written consent was provided, and participants received a taxable compensation of DKK 300. The study was approved by The Central Denmark Regional Committee on Health Research Ethics (case 1-10-72-11-15).

### Stimuli

Stimuli for the experiment were constructed from standard and deviant tones derived from Wizoo Acoustic Piano samples from Halion in Cubase 7 (Steinberg Media Technologies GmbH) with a 200 ms duration including a 5 ms rise time and 10 ms fall time.

Deviants differed from standards on one or more of three acoustic features: fundamental frequency (in Hz), sound intensity (in dB), and inter-aural time difference (in µs), henceforth referred to as *frequency* (F), *intensity* (I), and *location* (L), respectively. There are several reasons for focusing on these specific features. First, unlike alternative features such as duration (Czigler and Winkler 1996) and frequency slide (Winkler, Czigler, Jaramillo and Paavilainen 1998), the point of deviance can be established unambiguously for frequency, intensity, and location deviants. Second, thee features have reliably evoked additive MMN(m) responses in previous research; specifically, additivity has been demonstrated for frequency and intensity (e.g., Paavilainen et al. 2001) and frequency and location (e.g., Schröger 1995), but not yet for location and intensity or for all three features together. Finally, the restriction to three features balances representability and generalisability with practical feasibility within the typical timeframe of an MEG experiment.

In total, seven deviant types, comprising three single deviants, three double deviants, and one triple deviant, were generated through modification in Adobe Audition v. 3.0 (Adobe Systems Inc.). Specifically, frequency deviants (F) were shifted down 35 cents using the *Stretch* function configured to “high precision”. Intensity deviants (I) were decreased by 12 dB in both left and right channels using the *Amplify* function. Location deviants (L) resulted from delaying the right stereo track by 200µs compared to the left one. These parameter values were found to produce robust and relatively similar ERP amplitudes in a previous EEG study (Vuust, Liikala, Näätänen, Brattico and Brattico 2016). Double and triple deviants combining deviants in frequency & intensity (FI), intensity & location (IL), location & frequency (LF), and frequency & intensity & location (FIL) were generated by applying two or three of the operations just described, always in the order of frequency, intensity, and location.

### Procedure

Two MMN paradigms were used in the experiment. Specifically, as shown in Figure 3, four blocks (M1-M4) of the main complex paradigm, based on the musical multi-feature paradigm (cf., Vuust et al. 2011), were interleaved with three blocks (C1-C3) of a simpler control paradigm (cf., Näätänen, Pakarinen, Rinne and Takegata 2004) with lower degrees of musical relevance.

**Fig. 3.**
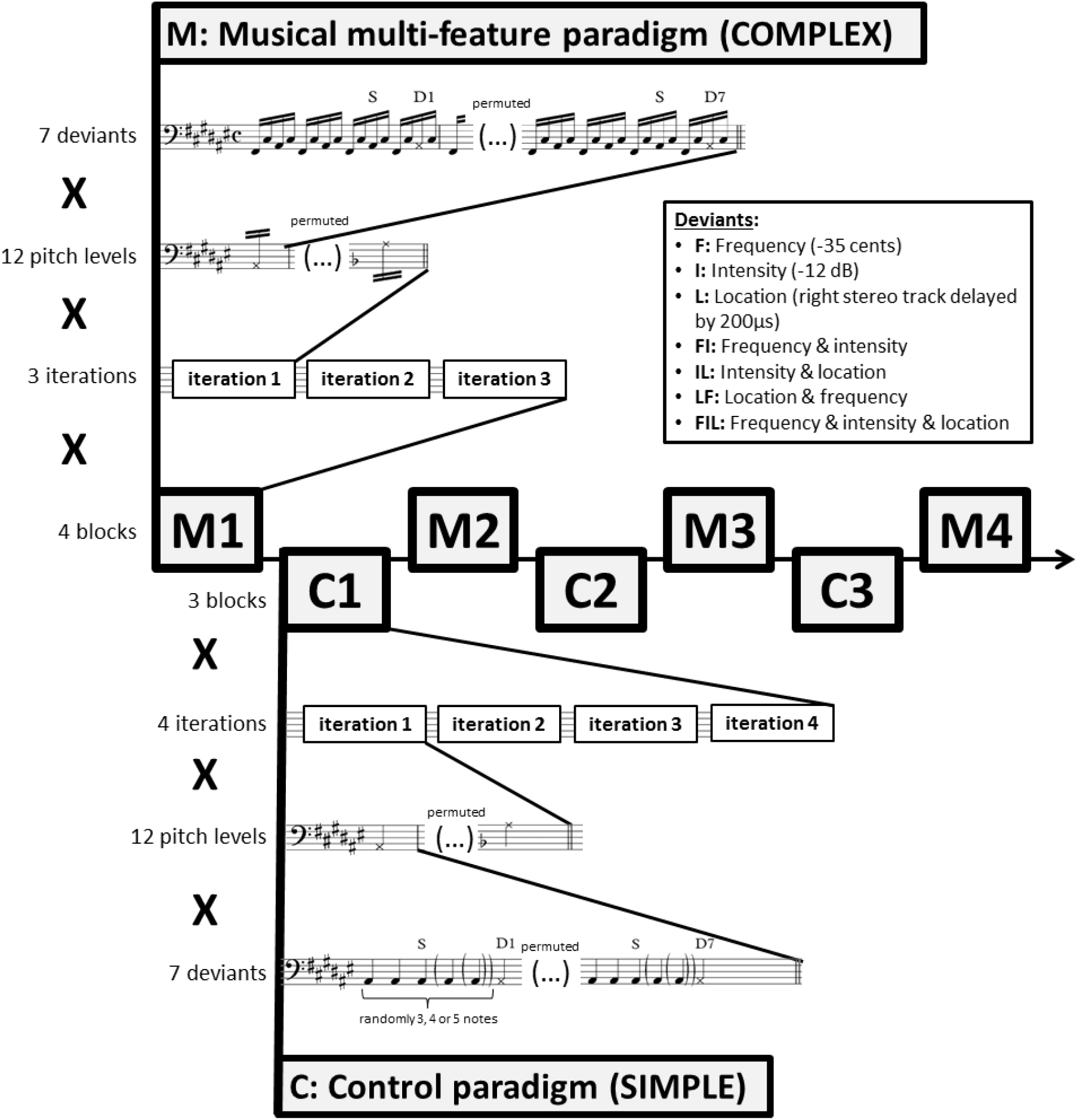
Experimental procedure. Stimuli were presented in alternating blocks using the complex musical multi-feature paradigm (M1-M4) and the simple control paradigm (C1-C3). Each block comprised three (M) or four (C) iterations of a sequence of standards and deviants presented at twelve varying pitch levels (i.e. chromatic range of A#2-A3). At each pitch level, seven deviant types were used in permutation. These comprised three single deviants differing in frequency (F, −35 cents), intensity (I, −12 dB), or location (L, right stereo track delayed by 200μs), double and triple deviants differing in frequency & intensity (FI), intensity & location (IL), location & frequency (LF), or frequency & intensity & location (FIL). Epoching of the magnetoencephalographic recordings was time-locked to the onset of the musical notes marked with S for standards and D for deviants.

The complex paradigm consists of repetitions of a characteristic four-note pattern referred to as the Alberti bass. In this pattern, the notes of a chord are arpeggiated in the order “lowest-highest-middle-highest” (see Figure 3). Although named after an Italian composer who used it extensively in early 18th-century keyboard accompaniment, the Alberti bass occurs widely across historical periods, instruments, and musical genres (Fuller 2015).

The studies introducing this paradigm (Vuust et al. 2011, 2016; Vuust, Brattico, Seppänen, Näätänen and Tervaniemi 2012a, 2012b; Timm et al. 2014; Petersen et al. 2015) have typically modified every second occurrence of the third note in the pattern (termed “middle” above) by changing its frequency, intensity, perceived location, timbre, timing, or by introducing frequency slides. In the present implementation, two extra occurrences of the standard pattern were introduced between each deviant pattern (see Figure 3) to minimize spill-over effects from consecutive deviants some of which made use of the same constituent deviant types due to the inclusion of double and triple deviants. Independence of deviant types is an underlying assumption of multi-feature mismatch negativity paradigms (Näätänen et al. 2004).

Consistent with previous studies, individual notes of the pattern were presented with a constant stimulus onset asynchrony (SOA) of 205 ms. After each occurrence of all seven deviants, the pitch height of the pattern changed pseudo-randomly across all 12 notes of the chromatic scale. This resulted in standard tones with frequency values ranging from 116.54 Hz (A#2) to 220.00 Hz (A3). Each block of the musical multi-feature paradigm comprised three such iterations of the 12 chromatic notes (Figure 3), resulting in a total of 144 trials of each deviant type across the four blocks.

The same number of trials per deviant type was obtained across the three blocks of the simple control paradigm, inspired by Näätänen, Pakarinen, Rinne and Takegata (2004). This was achieved by incorporating four rather than three iterations of the 12 chromatic notes (A#2 to A3) for each deviant array in each block (Figure 3). Instead of the Alberti bass, the simple paradigm used standard and deviant notes presented with a constant SOA of 400 ms, corresponding to a classical oddball paradigm. This was a multi-feature paradigm in the sense that it incorporated all seven deviant types in each block; however, the number of standards presented before each deviant varied randomly between 3 and 5, again to minimize spill-over effects from consecutive deviants with the same constituent deviant types (Figure 1).

During the complete experimental procedure (∼100 mins), participants were seated in a magnetically shielded room watching a silent movie with the soundtrack muted. Stimulus sounds were presented binaurally through Etymotic ER•2 insert earphones using the Presentation software (Neurobehavioral Systems, San Francisco, USA). Sound pressure level was set to 50 dB above individual hearing threshold as determined by a staircase procedure implemented in PsychoPy (Peirce 2007). Participants were instructed to stay still while the sounds were playing, to ignore them, and to focus on the movie. Complex musical multi-feature blocks (M1-M4) lasted 13 mins 47 sec whereas simple control blocks (C1-C3) lasted ∼11 mins. Between each block, short breaks of ∼1-2 mins were provided during which participants could stretch and move slightly while staying seated. Prior to the lab session, participants completed an online questionnaire that ensured eligibility and assessed their level of musical experience using *Goldsmiths Musical Sophistication Index*, v.1.0 (Müllensiefen et al. 2014).

### MEG recording and pre-processing

MEG data were sampled at 1000 Hz (with a passband of 0.03-330 Hz) using the Elekta Neuromag TRIUX system hosted by the MINDLab Core Experimental Facility at Aarhus University Hospital. This MEG system contains 102 magnetometers and 204 planar gradiometers. Head position was recorded continuously throughout the experimental session using four head position indicator coils (cHPI). Additionally, vertical and horizontal EOG as well as ECG recordings were obtained using bipolar surface electrodes positioned above and below the right eye, at the outer canthi of both eyes, and on the left clavicle and right rib.

Data were pre-processed using the temporal extension of the signal space separation (tSSS) technique (Taulu, Kajola and Simola 2004; Taulu and Simola 2006) implemented in Elekta’s MaxFilter software (Version 2.2.15). This included head movement compensation using cHPI, removing noise from electromagnetic sources outside the head, and down-sampling by a factor of 4 to 250 Hz. EOG and ECG artefacts were removed with independent component analysis (ICA) using the *find_bads_eog* and *find_bads_ecg* algorithms in MNE Python (Gramfort et al. 2013, 2014). These algorithms detect artefactual components based on either the Pearson correlation between the identified ICA components and the EOG/ECG channels (for EOG) or the significance value from a Kuiper’s test using cross-trial phase statistics (Dammers et al. 2008) (for ECG). Topographies and averaged epochs for the EOG- and ECG-related components were visually inspected for all participants to ensure the validity of rejected components.

Further pre-processing and statistical analysis were performed in FieldTrip (Oostenveld, Fries, Maris and Schoffelen 2011; http://www.ru.nl/neuroimaging/fieldtrip; RRID:SCR_004849). Data were epoched into trials of 500 ms duration including a 100 ms pre-stimulus interval. As indicated in Figure 1, for both paradigms, only the third occurrence of the standard tone after each deviant was included as a standard trial. Trials containing SQUID jumps were discarded using automatic artefact rejection with a *z* value cutoff of 30. The remaining 98.2% of trials on average (ranging from 91.0% to 99.8% for individual participants) were band-pass filtered at 1-40 Hz using a two-pass Butterworth filter (data-padded to 3 secs to avoid filter artefacts). Planar gradiometer pairs were combined by taking the root-mean-square of the two gradients at each sensor, resulting in a single positive value. Baseline correction was performed based on the 50 ms pre-stimulus interval.

### Experimental design and statistical analysis

To reiterate the experimental design, 25 musicians and 25 non-musicians completed a total of seven blocks distributed between the complex musical multi-feature paradigm (M) and simple control paradigm (C) in the order M1-C1-M2-C2-M3-C3-M4 (Figure 3). Trials were averaged for each condition (i.e. one standard and seven deviant types) separately for each participant and separately for each paradigm. Magnetic mismatch negativity responses (MMNm) were computed by subtracting the average standard (the third after each deviant, cf. Fig. 1) originating from the relevant paradigm from each of the deviant responses. These were the empirical MMNms. Consistent with previous studies (Levänen et al. 1993; Schröger 1995, 1996; Takegata et al. 1999; Paavilainen et al. 2001; Takegata, Wolff and Schröger, 2001; Paavilainen, Mikkonen, et al., 2003; Ylinen et al., 2005; Caclin et al., 2006; Lidji et al., 2009), modelled MMNms were computed for the three double deviants and one triple deviant by adding the two or three empirical MMNms obtained from the relevant single deviants.

The statistical analysis reported here focused on data from the combined planar gradiometers. Non-parametric, cluster-based permutation statistics (Maris and Oostenveld, 2007) were used to test the prediction that the additivity of the MMNm response would differ between musicians and non-musicians. Because violation of the additive model is assessed through demonstration of a difference between empirical and modelled MMNm responses, this hypothesis predicts an interaction effect between degree of additivity and musical expertise. This was tested by running cluster-based permutation tests on the difference between empirical and modelled MMNm responses, comparing between musicians and non-musicians. This approach for testing interaction effects within the non-parametric permutation framework is advocated by FieldTrip (see http://www.fieldtriptoolbox.org/faq/how_can_i_test_an_interaction_effect_using_cluster-based_permutation_tests).

The test statistic was computed in the following way: Independent-samples *t*-statistics were computed for all timepoint-by-sensor samples. Samples that were neighbours in time and/or space and exceeded the pre-determined alpha level of .05 were included in the same cluster. Despite its similarity to uncorrected mass-univariate testing, this step does not represent hypothesis testing, but only serves the purpose of cluster definition. To this end, a neighbourhood structure was generated that defined which sensors were considered neighbours based on the linear distance between sensors. Only assemblies containing minimum two neighbour sensors were regarded as clusters. A cluster-level statistic was computed by summing the *t* statistics within each cluster and taking the maximum value of these summed *t* statistics. This process was subsequently repeated for 10,000 random permutations of the group labels (musicians vs. non-musicians) giving rise to a Monte Carlo approximation of the null distribution. The final *p* value resulted from comparing the initial test statistic with this distribution. Bonferroni correction was applied to the two-sided alpha level to correct for the four comparisons of modelled and empirical double and triple MMNms, resulting in an alpha level of α = .025/4 = .00625. When significant additivity-by-expertise interactions were discovered, simple effects of additivity were assessed for musicians and non-musicians separately.

Previous research shows that the MMNm usually peaks in the 150-250 ms post-stimulus time range and is maximally detected with gradiometers bilaterally at supratemporal sites (Levänen et al. 1996; Näätänen et al. 2007). This was confirmed for the current dataset by computing the grand-average MMNm across all participants, all conditions, and both paradigms. This grand-average MMNm peaked in the combined gradiometer sensors MEG1342+1343 (right) and MEG0232+0233 (left) at ∼156 ms post-stimulus (with secondary peaks extending into the 200-300 ms range). However, to account for possible differences in peak latency and source location between participants and between the various deviant types, the analysis was extended to the 100-300 ms post-stimulus interval and to also include the eight neighbouring sensors around the peak sensor in each hemisphere (i.e. 18 sensors in total). In this way, prior knowledge was incorporated to increase the sensitivity of the statistical tests without compromising their validity (Maris and Oostenveld 2007).

Before the main analysis, however, two sets of tests were conducted to ensure the validity of the main analysis. To this end, cluster-based permutation tests were run using the parameters, sensors, and time window specified above (except for the permuted labels, which were changed according to the contrast of interest). First, the difference between standard and deviant responses was assessed to determine whether MMNm effects were indeed present, i.e. whether standard and deviant responses differed significantly in the 100-300 ms post-stimulus range (Table 1). These analyses were carried out for each deviant type separately for musicians and non-musicians and separately for the two paradigms. Second, to establish that the possible MMNm effects were potentially additive, nine pairwise comparisons were conducted between relevant single and double deviants as well as between relevant double and triple deviants (i.e., FI-FIL, IL-FIL, LF-FIL, F-FI, I-FI, I-IL, L-IL, L-LF, F-LF) (Table 2). This was done across all participants regardless of expertise level. No Bonferroni correction was applied to these secondary validity checks.

## Acknowledgements

The authors would like to thank Rebeka Bodak, Kira Vibe Jespersen, and the staff at MINDLab Core Experimental Facility, Aarhus University Hospital, for highly appreciated help with data collection and technical assistance.

